# Combinatorial effects on gene expression at the *Lbx1*/*Fgf8* locus resolve Split-Hand/Foot Malformation type 3

**DOI:** 10.1101/2022.02.09.479724

**Authors:** Giulia Cova, Juliane Glaser, Robert Schöpflin, Salaheddine Ali, Cesar Augusto Prada-Medina, Martin Franke, Rita Falcone, Miriam Federer, Emanuela Ponzi, Romina Ficarella, Francesca Novara, Lars Wittler, Bernd Timmermann, Mattia Gentile, Orsetta Zuffardi, Malte Spielmann, Stefan Mundlos

## Abstract

Split-Hand/Foot Malformation type 3 (SHFM3) is a congenital limb malformation associated with tandem duplications at the *LBX1*/*FGF8* locus. Yet, the disease patho-mechanism remains unsolved. Here we investigated the functional consequences of SHFM3-associated rearrangements on chromatin conformation and gene expression *in vivo* in transgenic mice. We show that the *Lbx1*/*Fgf8* locus consists of two separate, but interacting, regulatory domains. Re-engineering of a SHFM3-associated duplication and a newly reported inversion in mice resulted in restructuring of the chromatin architecture. This led to an ectopic activation of the *Lbx1* and *Btrc* genes in the apical ectodermal ridge (AER) in an *Fgf8-*like pattern. Artificial repositioning of the AER-specific enhancers of *Fgf8* was sufficient to induce misexpression of *Lbx1* and *Btrc*. We provide evidence that the SHFM3 phenotype is the result of a combinatorial effect on gene misexpression and dosage in the developing limb. Our results reveal new insights into the molecular mechanism underlying SHFM3 and provide novel conceptual framework for how genomic rearrangements can cause gene misexpression and disease.

## Introduction

At the sub-megabase scale, chromosomes are organized into distinct regions of high interaction called Topologically Associating Domains (TADs) that are separated from each other by boundaries^1,2^. TADs are thought to restrict enhancer-promoter contacts thereby defining regulatory domains. Structural variations (SVs), such as deletions, duplications and inversions, can disrupt these functional units and different types of SVs can induce distinct topological changes. For instance, deletions are mainly responsible for TAD fusion, duplications can promote formation of a new TAD (neo-TAD), while TAD shuffling is frequently observed upon an inversion^3–5^. SVs can not only affect gene dosage but also modulate basic mechanisms of gene regulation. Indeed, SVs can alter the copy number of regulatory elements or modify the 3D genome by disrupting higher-order chromatin organisation. This can give rise to ectopic enhancer-promoter contacts, gene misregulation and ultimately disease^5^. However, the general applicability of this concept remains under investigation. Studies in Drosophila, for example, demonstrated that the disruption of TADs and enhancer-promoter interactions are not necessarily accompanied by changes in expression, indicating a high degree of robustness of the genome against such events6.

SHFM is a congenital limb malformation characterized by variable defects of the central rays of the autopod often together with syndactyly and/or aplasia/hypoplasia of the phalanges, metacarpals, and metatarsals^7^. SHFM is known as a paradigm for genetic heterogeneity. While duplications on either 17p13.3 or 10q24^8^ are the most frequent genetic cause, SHFM can be also associated with coding and non-coding mutations at other loci. Indeed, SVs involving the *DYNC1I1* locus where regulatory elements for *DLX5* and *DLX6* are localized^9^, and mutations in *TP63*, *DLX5*, or *WNT10B*^10^ have been previously associated with SHFM. The common underlying patho- mechanism of these mutations is thought to be the failure to maintain a proliferation/growth signal from the apical ectodermal ridge (AER), a signalling centre of ectodermal cells at the distal end of the limb bud^11^. SHFM is also a clinically heterogeneous disease^12^, as some patients carrying SHFM-associated SVs do not show any limb phenotype or a very mild one, suggesting that SHFM patho-mechanism is not merely associated to gene dosage alteration.

Tandem duplications on chromosome 10q24 at the *LBX1*/*FGF8* locus are considered the cause for SHFM type 3 (SHFM3)^13^, but the specific patho-mechanism of SHFM3 associated duplications remains unsolved. The reported duplications are at least 500 kb in size, encompassing five genes of the locus (*LBX1*, *BTRC*, *POLL*, *DPCD*, and *FBXW4*) but excluding the neighbouring *FGF8* (**Supplementary Fig. 1**)^14^. While *LBX1* is essential for early muscle cell differentiation and migration^15^, *FGF8* is expressed in the AER together with other FGFs where it functions as a growth factor for the underlying mesechyme^16^. *BTRC* is important in regulating cell cycle checkpoints^17^, *DPCD* in the generation and maintenance of ciliated cells^18^, *POLL* encodes for a DNA polymerase^19^, whereas *FBXW4* is involved in ubiquitin mediated protein degradation^20^. Duplications can have different effects on gene regulation depending on their relative position to chromatin domain boundaries. Studies of the *SOX9* locus, for example, have shown that duplications within the *SOX9* TAD can lead to *SOX9* mis/overexpression and sex reversal, whereas duplications that include the *SOX9/KCNJ2* boundary and the *KCNJ2* gene result in *KCNJ2* misexpression and brachydactyly^4^. In the latter situation, a neo-TAD is formed leading to contact of the *SOX9* enhancers with the *KCNJ2* promoter and *KCNJ2* expression in a *SOX9-like* pattern.

Given these examples, it is conceivable that the SHFM3-associated duplications may have specific regulatory effects that could result in gene misexpression.

In addition to the SHFM3 duplications at the *LBX1*/*FGF8* locus, we reported here for the first time a new case of SHFM3 malformation associated with an inversion at the same locus. This inversion was also of particular interest as the patient is a good example of clinical heterogeneity of this limb malformation within the same individual (**Supplementary Fig. 2**). In order to uncover the molecular mechanism underlying this congenital disease, we engineered a duplication and the inversion at the *Lbx1/Fgf8* locus in mice using CRISPR/Cas9 genome editing tool^21^. Chromosome conformation capture Hi-C (cHi-C) analysis of the mouse developing limbs in mutants showed that both SVs led to a disruption of the wild-type three-dimensional chromatin configuration of the locus and ectopic enhancer-promoter contacts. Strikingly, this caused misexpression of two genes, *Lbx1* and *Btrc*, in an *Fgf8*-like pattern in the AER which appeared to be associated with a dose-dependent phenotype. Collectively, our results illustrate the impact of SVs at the *Lbx1/Fgf8* locus on chromatin architecture and gene regulation and reveal the complex molecular patho- mechanism underlying SHFM3 limb malformation. With the emergence of novel sequencing technologies and the improvement in the discovery and characterization of SVs, this study illustrates the myriad ways by which SVs can cause disease (i.e., a combination of enhancer repositioning and dosage effect), highlighting the challenging task of their interpretation in the clinic.

## Results

### SHFM3-associated SVs include the *Lbx1*/*Fgf8* domain boundary and the *Fgf8* AER enhancers

To characterize the 3D conformation at the *Lbx1*/*Fgf8* locus during mouse development, we performed cHi-C in wild-type murine limb buds at E11.5, a developmental stage where both *Lbx1* and *Fgf8* limb developmental genes are expressed (**Fig. 1a**). We observed that the locus is characterized by the presence of two main chromatin regulatory domains (**Fig. 1a,** indicated by black dashed lines), a centromeric and a telomeric containing *Lbx1/Btrc* and *Fbxw4*/*Fgf8* genes, respectively, and hereafter referred to as *Lbx1* and *Fgf8* TADs. These domains are separated from neighbouring TADs and from each other by CTCF-associated TAD boundaries in the classical divergent orientation^22^ (**Fig. 1a**). *Poll* and *Dpcd* are located within the boundary region separating *Lbx1* and *Fgf8* TADs. Given the divergent roles and expression pattern of *Lbx1* and *Fgf8* during muscle and limb development, respectively^23,24^, separated regulatory domains can be expected to ensure correct gene regulation. The *Fgf8* TAD contains several well characterized enhancers with AER-specific activity as well as other tissue-specific enhancers corresponding to the *Fgf8* expression pattern in the embryo, such as the branchial arches, the somites and the midbrain-hindbrain boundary^25,26^. Interestingly, all identified AER enhancers (58, 59, 61, 66; yellow ovals in **Fig. 1a**) are located within a 40 kb region in the introns of *Fbxw4*, with the exception of one enhancer (80; orange oval in **Fig. 1a**) which is located in close proximity to *Fgf8*.

**Fig. 1.**
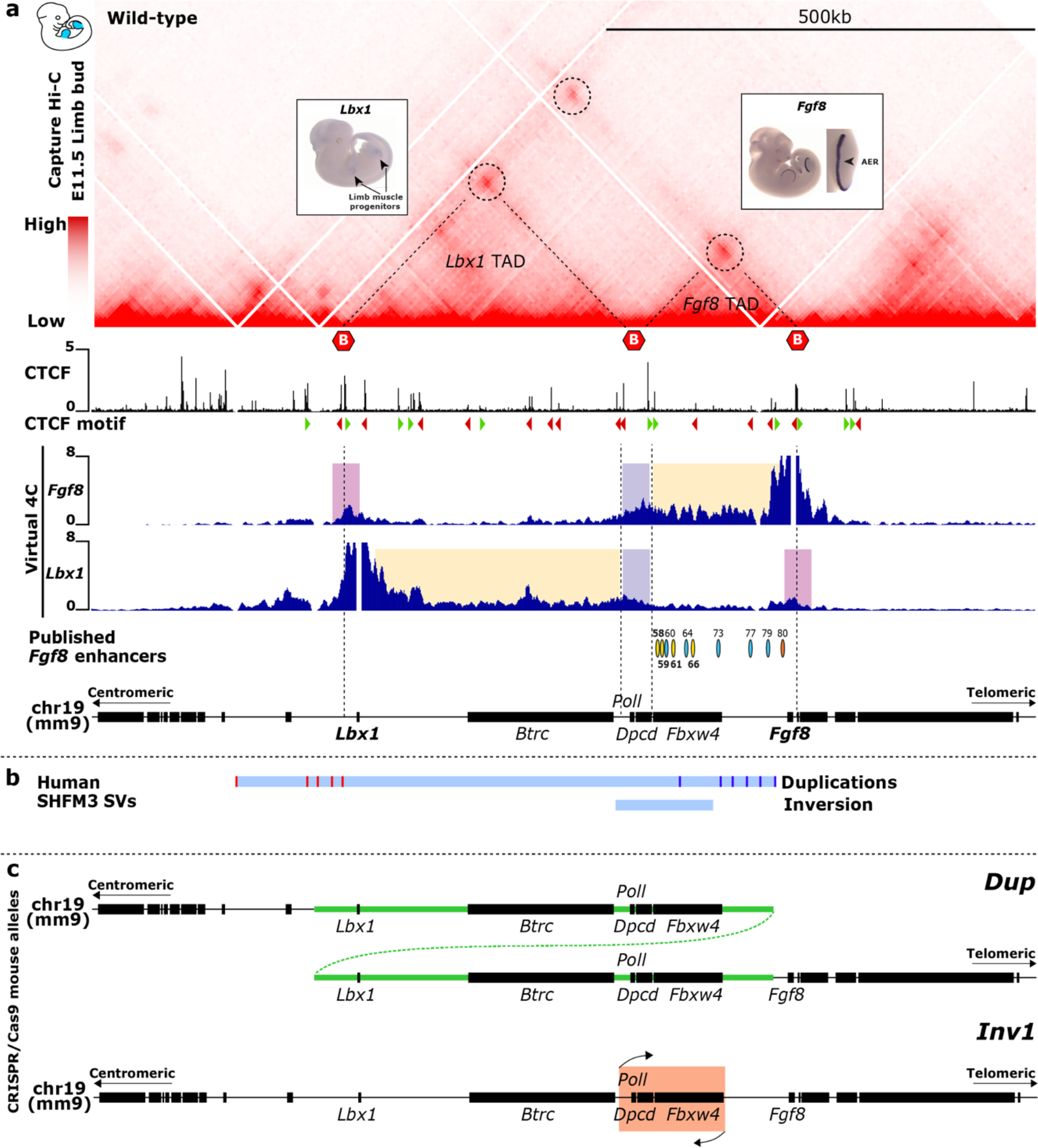
3D chromatin architecture at the *Lbx1*/*Fgf8* locus and localisation of SHFM-associated SVs. **a**, cHi-C of extended *Lbx1*/*Fgf8* locus generated from wild-type E11.5 mouse limb buds. The region consists of two TADs (indicated by dashed lines) separated by boundaries (indicated by red hexagons) in correlation with CTCF binding sites, as indicated in ChIP-seq from E11.5 mouse limb buds^54^ below. Loops (dashed circles) indicate interaction between CTCF sites. Interaction plots using virtual 4C from *Fgf*8 and *Lbx1* viewpoints are shown below. Regions of interactions relative to *Lbx1* and *Fgf8* within their own TADs are highlighted in yellow, whereas contacts with the boundary region between *Lbx1* and *Fgf8* TADs and those over such boundary are in violet and pink, respectively. Published *Fgf8* enhancers25 are indicated by ovals. Yellow ovals highlight enhancers driving *Fgf8* expression in the AER and localized within the introns of *Fbxw4*, while in orange is the only AER enhancer in close proximity to *Fgf8*. **b,** Schematic of human SHFM3 related structural variations (SVs). Red and blue lines in the duplications bar represent the different centromeric and telomeric breakpoints, respectively. Breakpoints of the inversion are shown below. **c,** Schematic of the SHFM-associated SVs re-engineered in mice using CRISPR/Cas9 genome editing tool, particularly one selected tandem duplication (*Dup*) and the inversion (*Inv1*).

Virtual 4C analysis with viewpoints on *Lbx1* and *Fgf8* further emphasized how the interactions of these genes are mainly restricted within their respective chromatin regulatory domains (interactions are highlighted in yellow in **Fig. 1a**). As *Fgf8* and *Lbx1* are located very close to the centromeric and telomeric boundaries, respectively, in the virtual 4C we could also appreciate the interactions with the TAD boundary region (highlighted in violet in **Fig. 1a**) defining the two domains. Despite being two separated domains, the *Lbx1* and *Fgf8* TADs are also connected via an additional loop (**Fig. 1a**, top dashed circle in cHi-C map). This interaction across the boundary is also clearly visible in our 4C analysis (highlighted in pink in **Fig. 1a**) indicating that subpopulations of limb bud cells can have different configurations, but also that a certain degree of leakiness of the *Lbx1/Fgf8* boundary exists, leading to a certain level of intermingling of the two domains in spite of the very different expression patterns of *Lbx1* and *Fgf8*.

Mapping of in-house and already published SHFM3 duplications onto the 3D structure of the locus showed that they all include the boundary between the *Lbx1* and *Fgf8* TADs (**Fig. 1b**). Furthermore, all duplications comprised *Lbx1* at the centromeric side and the previously identified *Fgf8* AER enhancers located within *Fbxw4*^26^, while excluding *Fgf8* at the telomeric side (**Fig. 1b and Supplementary Fig. 1**). Upon mapping of our new case of SHFM inversion we observed that the boundary region between the *Lbx1* and *Fgf8* TADs and the *Fgf8* AER enhancers are involved also in this SHFM-associated SV, suggesting a potential common patho-mechanism.

### SHFM3-associated duplication and inversion result in ectopic interactions of *Fgf8* AER enhancers region with *Lbx1* TAD

To investigate the effect of the SHFM3-associated duplications and inversion on chromatin configuration, gene expression and phenotype, we engineered a human SHFM3 duplication (*Dup*) and the newly reported inversion (*Inv1*) in mice (**Fig. 1c**). cHi-C was performed in homozygous *Dup* and *Inv1* E11.5 murine limb buds (**Fig. 2**). The *Dup* cHi-C map (**Fig. 2b**, LOIC-normalized) showed a general increase in the frequency of interactions within the duplicated DNA region compared to the wild-type (**Fig. 2a**), reflecting the copy number increase. Interestingly, in the subtraction of scaled KR-normalized maps (*Dup* versus wild-type), in which the effect of the higher copy number on the signal intensity is reduced (**Fig. 2c**), we observed a stripe of increased interactions (indicated by black arrowheads in **Fig. 2c**) between part of the *Fgf8* TAD containing the *Fgf8* AER enhancers involved in the duplication and a region immediately upstream of the *Lbx1* centromeric TAD boundary. This reflected the formation of a neo-TAD, as illustrated in the schematic (**Fig 2d**). The neo-TAD, indicated by the blue/green triangle between the two *Lbx1* TADs (yellow), contained only *Fbxw4*, the *Fgf8* AER enhancers and a small (∼38 kb) region flanking the centromeric side the *Lbx1* TAD boundary, thus unlikely to have a pathological effect by itself. However, we also observed increased contacts between the entire *Lbx1* TAD and the *Fgf8* AER enhancers region (indicated by dashed rectangle and asterisks in **Fig. 2b and c**). These interactions were likely established by the *Fgf8* AER enhancers from the neo-TAD, with both the endogenous and duplicated *Lbx1* TADs (red arrows in **Fig. 2d**). Indeed, in the *Dup* mutant, interaction between the *Fgf8* AER enhancers and the 3’ *Lbx1* TAD is favoured by a shorter distance (the *Lbx1* gene is located ∼130kb in *Dup* versus ∼380kb in wild-type configuration) and by the presence of a weaker TAD boundary (two CTCF binding sites instead of four). Moreover, the *Fgf8* AER enhancers could also contact the 5’ *Lbx1* TAD due to the previously mentioned leakiness of the boundary and the absence of their target gene *Fgf8*, making these enhancers free to interact with the neighbouring *Lbx1* gene and the entire *Lbx1* TAD.

**Fig. 2.**
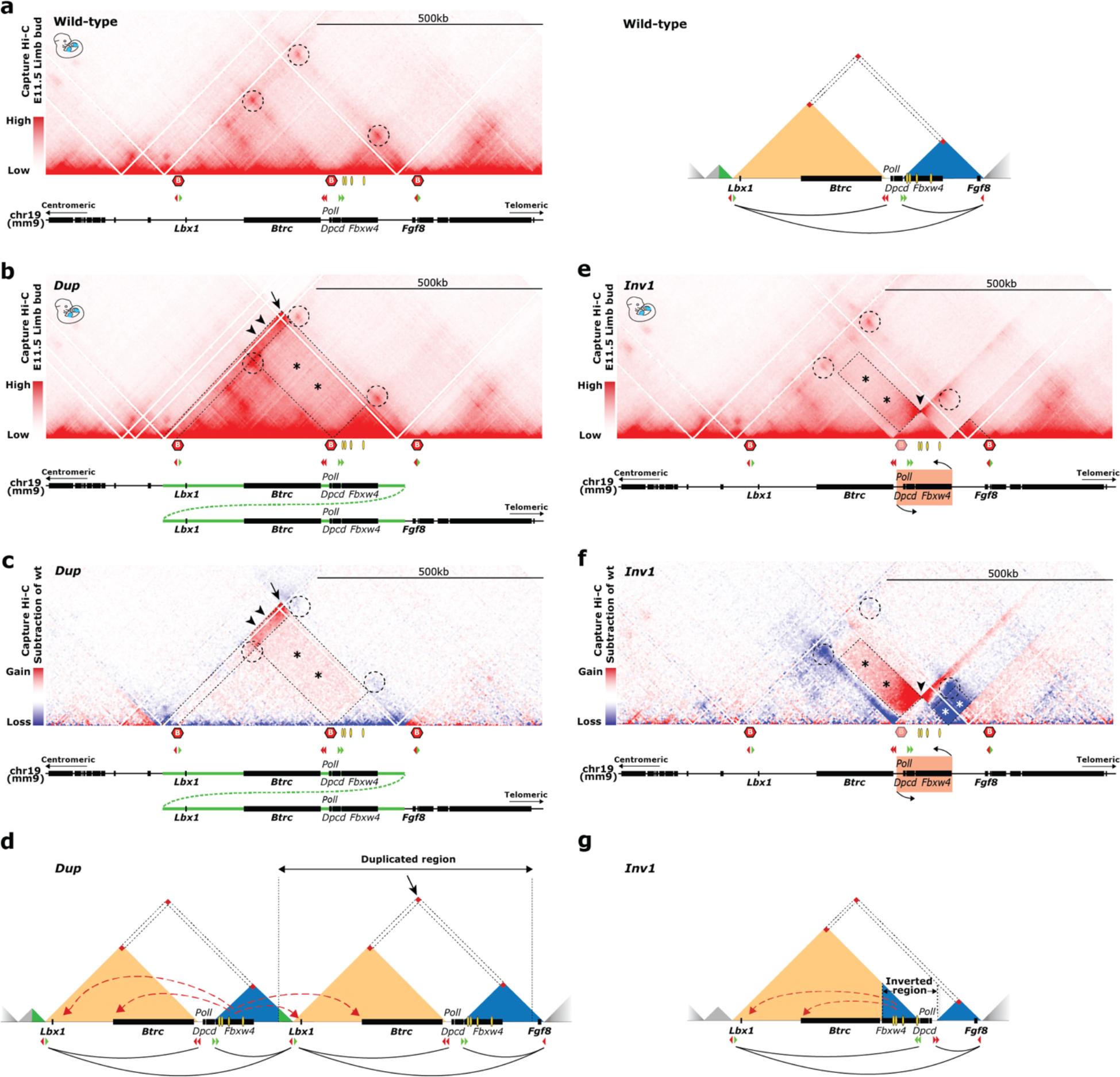
Ectopic interactions upon structural rearrangements at the *Lbx1*/*Fgf8* locus. **a**, Wild-type cHi-C from E11.5 mouse limb buds (left panel). Boundaries are indicated by red hexagons and interactions between boundaries are highlighted by dashed circles. Orientation of CTCF binding sites and schematic of the locus are shown below. Yellow ovals highlight the *Fgf8* AER enhancers. Schematic of wild-type configuration is shown on the right (right panel). *Lbx1* TAD in yellow, *Fgf8* TAD in blue. Interactions between boundaries are highlighted by red squares. Orientation of CTCF binding sites, interactions between CTCF in convergent orientation and schematic of the locus are shown below. **b,** cHi-C of homozygous *Dup*. Positions of preserved wild-type interactions between boundaries are indicated by dashed circles. The new contact reflecting the breakpoints of the duplicated region is highlighted by a black arrow. Black arrowheads indicate a strong gain of contacts between the region containing the *Fgf8* AER enhancers and the region immediately flanking the centromeric side of the *Lbx1* TAD boundary. The rectangular dashed area highlighted by asterisks show increased interactions between the *Fgf8* AER enhancers region and the *Lbx1* TAD. **c,** Subtraction of wild-type cHi-C interactions from the *Dup* mutant. The new contact reflecting the breakpoints of the duplicated region is highlighted by a black arrow. Central and telomeric TAD boundary interactions are not affected (central and telomeric dashed circles), while the centromeric dashed circle highlights a gain of interaction representative of the new extra *Lbx1* TAD. Black arrowheads and rectangular dashed area (asterisks) point to the increased interactions involving the *Fgf8* AER enhancers region. **d,** Schematic of *Dup* configuration. The entire *Lbx1* TAD is duplicated and remains unchanged in its content and configuration. The telomeric *Fgf8* TAD also remains unchanged. Between the two *Lbx1* TADs a neo-TAD forms containing the *Fbxw4* gene, the *Fgf8* AER enhancers and the small region centromeric of the *Lbx1* TAD boundary. Red dashed arrows indicate the ectopic contacts between *Fgf8* AER enhancers and the *Lbx1* TAD. **e,** cHi-C of homozygous *Inv1*. Dashed circles indicate the position of the original wild-type interactions between boundaries, two of them lost upon reshuffling of the boundary between *Lbx1* and *Fgf8* TADs as a consequence of the inversion. The boundary involved in the inversion is shown as blurry. Dashed lines on the right point out the new smaller *Fgf8* TAD, now comprising only *Fgf8*. Black arrowhead highlights the bow tie configuration representative of the inverted regions. The rectangular dashed area highlighted by asterisks show ectopic interactions compared to wild- type between the *Fgf8* AER enhancers region and the *Lbx1* TAD. **f,** Subtraction of wild-type cHi-C interactions from the *Inv1* mutant. Centromeric and telomeric dashed circles highlight the loss (blue) of the original interactions between boundaries as a consequence of TAD-reshuffling. Black arrowhead highlights the classic bow tie configuration representative of the inverted regions. The rectangular dashed area highlighted by black asterisks show ectopic interactions compared to wild-type between the *Fgf8* AER enhancers region and the *Lbx1* TAD, while the square dashed area highlighted by white asterisks points out the loss of interactions between *Fgf8* and its AER enhancers. **g,** Schematic of *Inv1* configuration. Upon inversion, the *Fgf8* AER enhancers are repositioned within a bigger *Lbx1* TAD and can now establish ectopic interactions (red dashed arrows), whereas the *Fgf8* TAD is reduced in size due to the inversion of the boundary and contains now only *Fgf8*, isolated from its AER enhancers.

We observed ectopic interactions between the *Fgf8* AER enhancers region and the *Lbx1* TAD also in *Inv1* cHi-C (**Fig. 2e and f**). In the inversion, the boundary between *Lbx1* and *Fgf8* TADs functioned as an anchor point around which part of the *Fgf8* TAD was moved into the *Lbx1* TAD and vice versa. Due to the fact that the centromeric breakpoint of the inversion was relatively close to the boundary, the *Lbx1* TAD was enlarged while the *Fgf8* TAD was reduced in size. As a consequence, the *Fgf8* AER enhancers were now relocated within this bigger *Lbx1* TAD which exhibited increased contacts (dashed rectangle and asterisks in **Fig. 2e and f**) with the repositioned *Fgf8* AER enhancer region, underlining a common trait with the *Dup*. Additionally, upon such reshuffling, *Fgf8* was now isolated from its own AER enhancers and we observed indeed loss of interactions between these elements and their endogenous gene (dashed rectangle with white asterisks in **Fig. 2f**).

Given that the overall configuration of the locus with two main TADs was conserved in humans **(Fig. 3a**), we then investigated whether the observed interaction changes were also present in SHFM3 patients. We performed 4C-seq experiments in human fibroblasts from a patient carrying the SHFM3 duplication (fibroblasts from the inversion case were not available) and exhibiting the classical SHFM phenotype. Using viewpoints on *FGF8* and on the two genes in the *LBX1* TAD (*LBX1* and *BTRC*), we detected ectopic interactions involving the *FGF8* AER enhancers region and both *LBX1* and *BTRC* which were not present in the healthy control (**Fig. 3b**). Comparison with mouse virtual 4C plots generated from *Dup* cHi-C maps further confirmed that the same ectopic interaction existed also in our *Dup* mutants (**Fig. 3c**), revealing we generated here a genetic mutant mouse model for SHFM3.

**Fig. 3.**
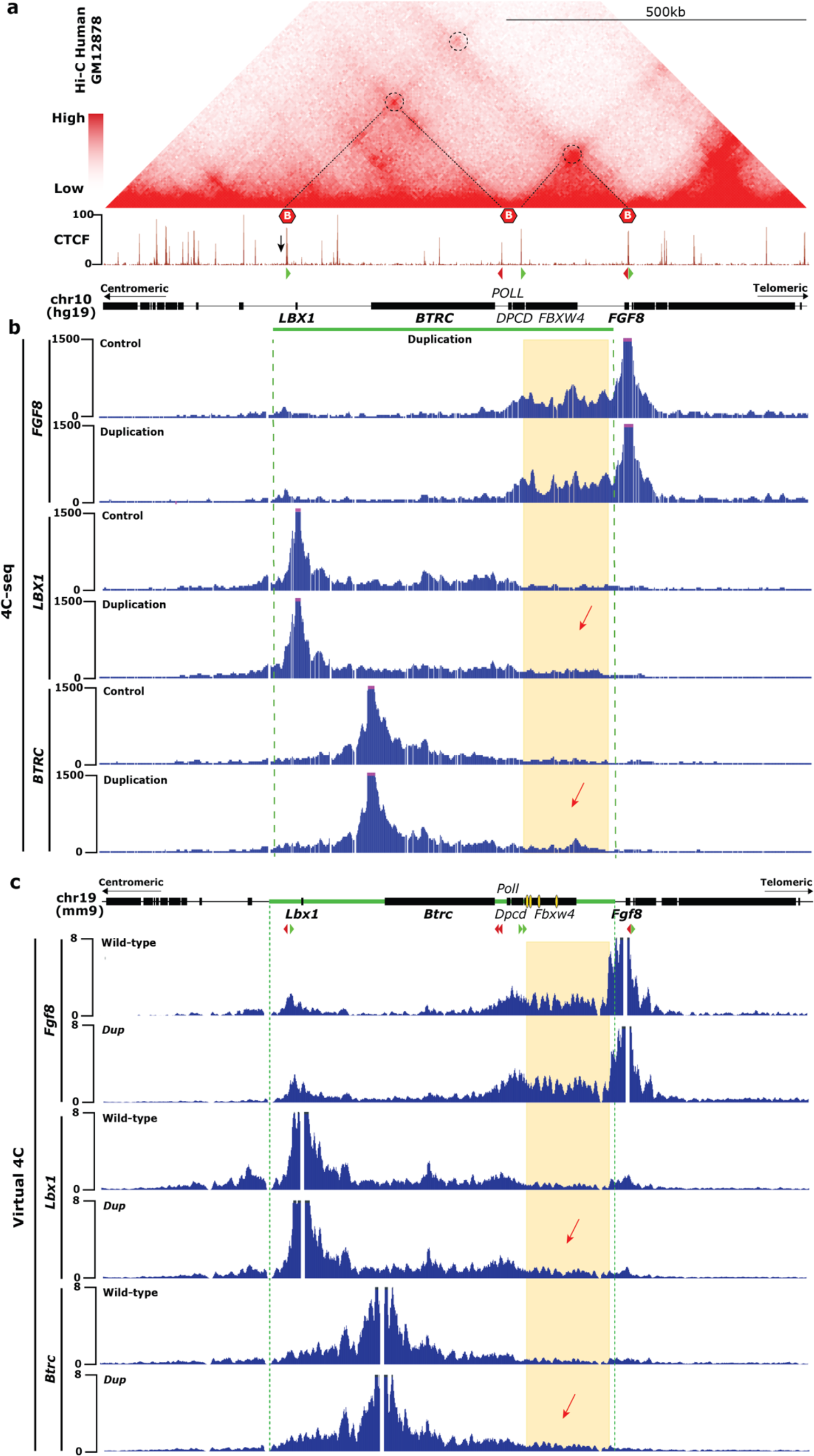
4C-seq in human fibroblasts reveals ectopic interactions involving *LBX1* and *BTRC*, further supported by virtual 4C in mouse. a, Hi-C at the *LBX1*/*FGF8* locus (hg19; chr10:102,669,128-103,840,922) derived from GM12878 cell^43^. TADs structures are similar between human and mouse. CTCF ChIP-seq from GM12878 cells^55,56^, orientation of CTCF binding sites and schematic of the human *FGF8* locus are shown below. Black arrow indicates a CTCF binding site in negative orientation at the *LBX1* centromeric TAD boundary present in mouse but not in human. b, 4C-seq in human fibroblasts from a healthy control and a patient carrying the SHFM duplication with viewpoints on *FGF8*, *LBX1* and *BTRC* promoter regions. The yellow area uncovers the region containing the *Fgf8* enhancers and the red arrows highlights the gain of ectopic interactions of this region with *LBX1* and *BTRC* upon duplication. Green bar and dashed lines indicate the duplicated region. C, Virtual 4C plots generated from cHi-c *Dup* maps with viewpoints on *Lbx1*, *Fgf8* and *Btrc* promoter regions. The yellow area uncovers the region containing the *Fgf8* enhancers (those specific for the AER are indicated as yellow ovals in the schematic of the murine locus) and the red arrows highlights the gain of ectopic interactions of this region with *Lbx1* and *Btrc* upon duplication. Green bar and dashed lines indicate the duplicated region.

### *Dup* and *Inv1* result in ectopic expression of *Lbx1* and *Btrc* in an *Fgf8*-like pattern in the AER

To investigate the effect of the observed chromatin rearrangements on gene expression, we performed detailed expression analysis (RNA-seq, single cell RNA-seq and whole mount *in situ* hybridisation (WISH)) at developmental stage E11.^5^. In the *Dup* mutant, all genes within the duplication (*Lbx1*, *Btrc*, *Poll*, *Dpcd* and *Fbxw4*) were significantly upregulated in the developing limb buds, corresponding to the copy number increase (**Fig. 4a**). In the *Inv1* mutant, *Fbxw4* and *Fgf8* were downregulated, reflecting the disruption of the gene and the repositioning of the *Fgf8* AER enhancers, respectively (**Fig. 4a**). Strikingly, *Lbx1* and *Btrc* exhibited increased expression levels in the *Inv1* mutant, indicating that the observed chromatin rearrangement had an effect on gene expression. Genes outside of the *Fgf8* and *Lbx1* TADs (*Sif2*, *Twnk* and *Oga*) remained unchanged in both *Dup* and *Inv1* mutants (**Fig. 4a**).

**Fig. 4.**
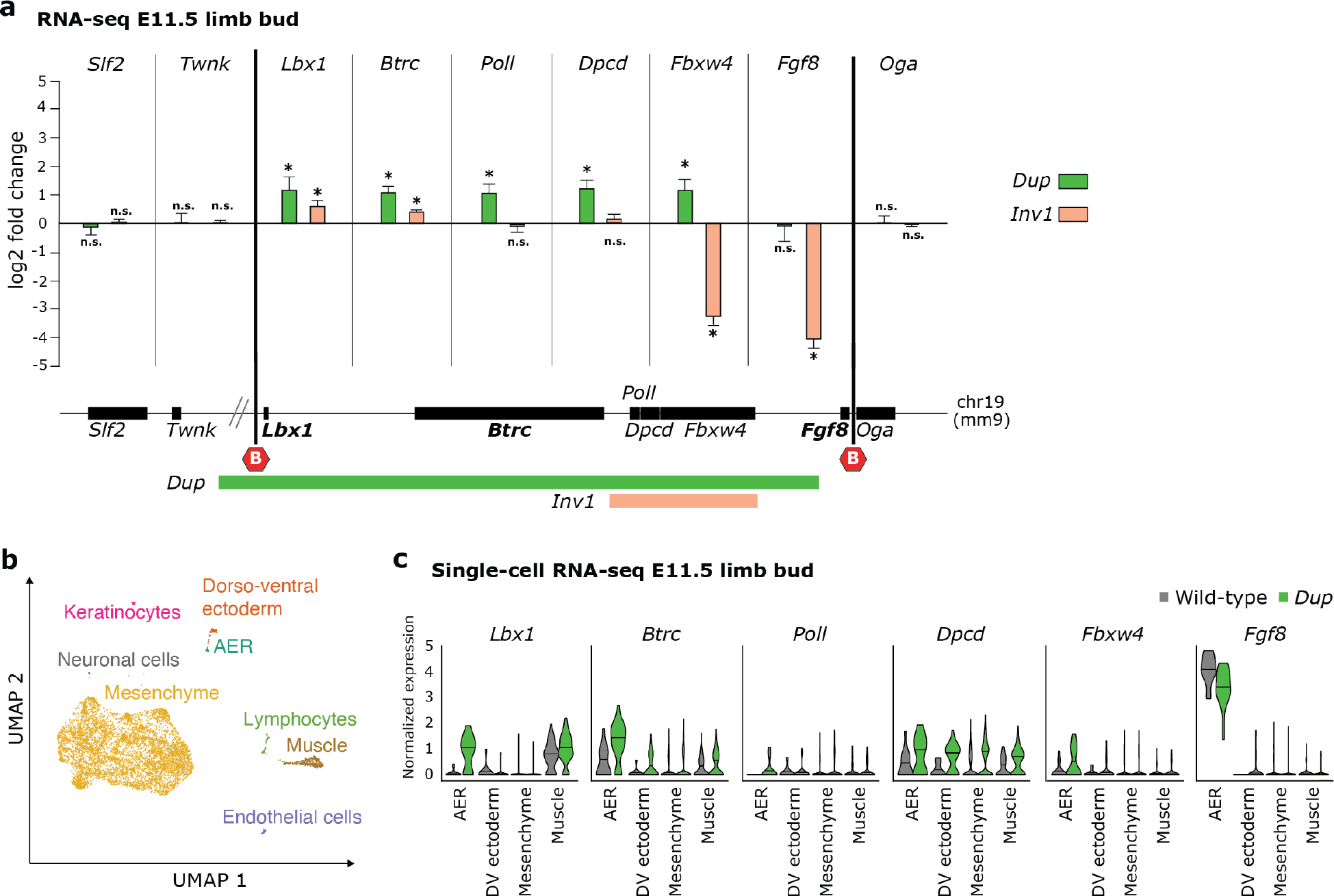
*Dup* and *Inv1* chromatin rearrangements have an impact on gene expression. **a**, Bar plot representing the log2 fold change expression compared to wild-type of genes in the *Lbx1* and *Fgf8* TADs and flanking genes (*Slf2* and *Twnk* (centromeric), *Oga* (telomeric)) from RNA-seq data of *Dup* and *Inv1* E11.5 limb buds. Schematic of the locus shows the position of the genes, the TAD boundaries (red hexagons) and the parts included in the *Dup* and the *Inv1* mutants. Wild-type *Dup* and *Inv1* samples are shown in black, green and orange, respectively. Error bars represent standard error, n.s = non-significant, *Adjusted Wald p-value < 0.05. **b,** Uniform manifold approximation and projection (UMAP) showing 8 cell clusters identified via scRNA-seq of E11.5 mouse hindlimbs from wild-type and *Dup* mutant. **c,** Violin plot representing the normalized expression in AER, dorso-ventral (DV) ectoderm, mesenchyme and muscle cells for the 6 genes that are part of the *Lbx1* and *Fgf8* TADs. Wild-type and *Dup* samples are shown in black and green respectively.

To investigate whether these changes of gene expression were specific to some regions of the limb bud, we performed single-cell RNA-seq (scRNA-seq) from E11.5 wild-type and *Dup* limb buds. Bioinformatic analysis revealed individual clusters corresponding to the major cell types in the developing limb including mesenchyme, dorso-ventral ectoderm, AER and muscle as well as satellite cell types such as neurons, lymphocytes, keratinocytes and endothelial cells (**Fig. 4b**). In wild-type cells, *Fgf8* expression was only present in the AER, whereas *Lbx1* was expressed only in muscle cells and the four other genes at the locus were expressed in all major cell types of the developing limb (**Fig. 4c**). The bulk RNA-seq data were confirmed such that the five genes included in the duplication (*Lbx1, Btrc, Poll, Dpcd and Fbxw4*), but not *Fgf8*, displayed increased expression in the *Dup* mutant compared to wt (**Fig. 4c**). Interestingly, *Lbx1*, and to a lesser extent *Btrc,* showed a strong AER expression in the *Dup* mutant, indicating misexpression in the AER on top of the increased copy number. This suggested that both *Lbx1* and *Btrc* were activated by the *Fgf8* AER enhancers now able to contact the two genes’ promoters. Thus, the scRNA-seq data confirmed the bulk RNA-seq data and showed, in addition, that *Lbx1* and *Btrc* were misexpressed in the AER.

Next, we performed WISH to confirm and further investigate potentially altered expression patterns. We uncovered that *Lbx1*, normally expressed in migrating muscle cells (black arrowheads for *Lbx1* in **Fig. 5a**) and *Btrc*, a ubiquitously expressed gene, were ectopically expressed in the AER, resembling the *Fgf8* specific expression pattern (**Fig. 5a**). As observed in the scRNA-seq data, *Lbx1* misexpression appeared to be stronger than *Btrc* (**Fig. 4c and Fig. 5a**). Importantly, an even more pronounced misexpression of both genes was also observed in the *Inv* mutant (**Fig. 5a**). In the *Dup* mutant, *Fgf8* expression was unchanged compared to wild-type. In contrast, in the *Inv1* mutant we observed a loss of *Fgf8* expression likely due to the loss of the endogenous interaction of *Fgf8* with its enhancers (**Fig. 5a**). Finally, we showed that misexpression in the AER was not detected for *Poll*, *Dpcd* and *Fbxw4*, neither in *Dup*, nor in the *Inv1* mutants (**Supplementary Fig. 3a**).

**Fig. 5.**
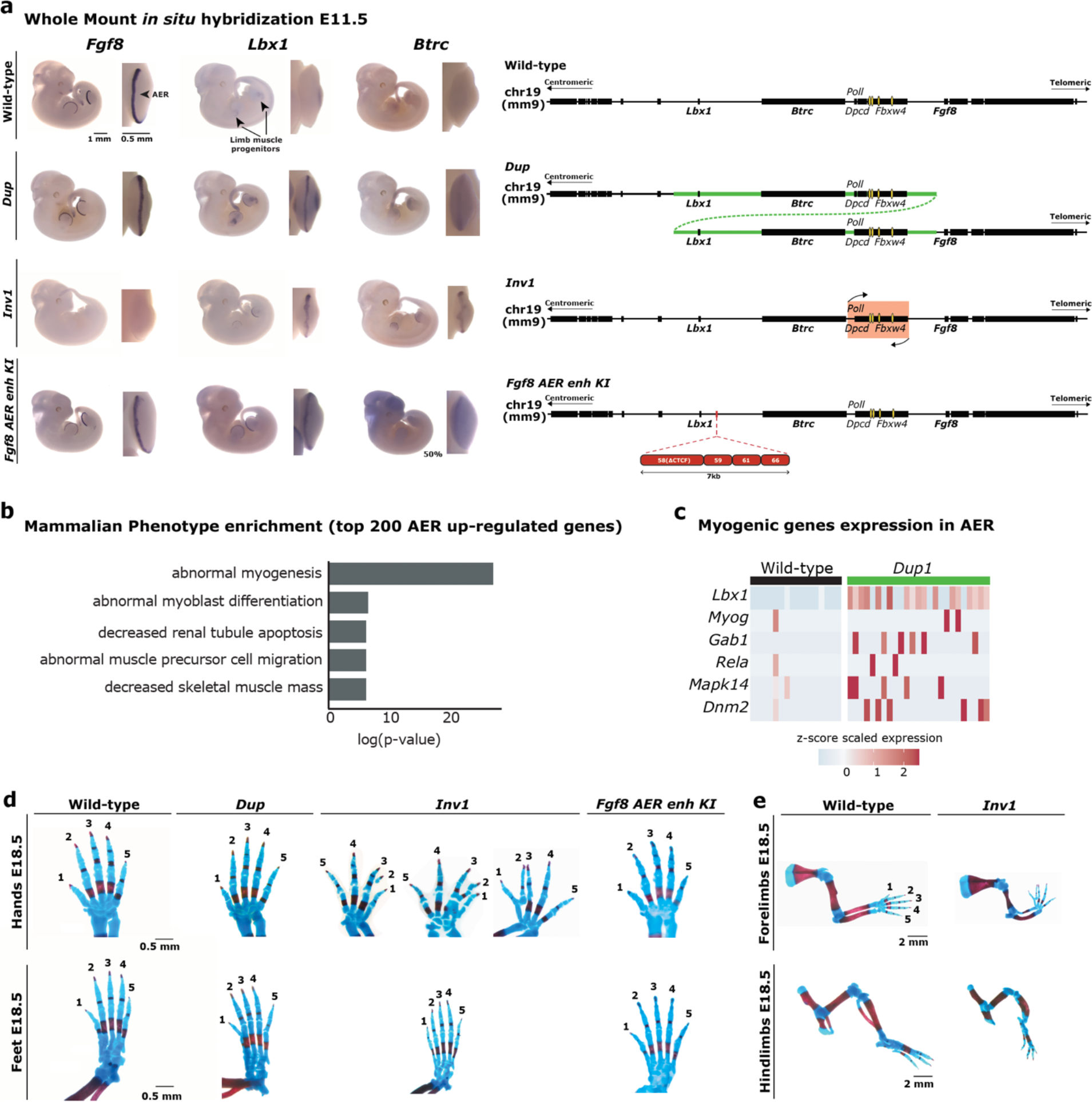
Ectopic interactions led to *Lbx1* and *Btrc* misexpression in the AER. **a**, Whole-mount *in situ* hybridization for *Fgf8*, *Lbx1* and *Btrc* at E11.5. Expression was checked and confirmed in at least 3 or more homozygous embryos (at least *n*=3 biological replicates) (left panel). Schematics of the locus for wild-type and CRISPR/Cas9 alleles (right panel). Yellow ovals highlight *Fgf8* AER enhancers. **b,** GO analysis for the MGI Mammalian phenotype enrichment terms using the top 200 genes that are up-regulated genes in the *Dup* mutant compared to wild-type in the AER cells from the scRNA-seq data. The 5 most significant enriched terms are represented on a log (p-value) scale. **c,** Heatmap showing the expression of 6 genes (*Lbx1, Myog, Gab1, Rela, Mapk14* and *Dnm2*) associated with the myogenic enriched terms in (**b**) in the AER cells. Each column represents one AER cell and the z-score scaled expression is indicated for each gene, showing a global increased expression in the *Dup* mutant’s AER cells. **d,** Skeletal analysis of E18.5 limbs stained with alcian blue (cartilage) and alizarin red (bone). Hands and feet from wild-type and homozygous *Dup*, *Inv1* and *Fgf8 AER enhancer KI* E18.5 embryos. No particular phenotype was observed in feet from all the mutants and in hands from *Dup* and *Fgf8 AER enhancer KI*. Fused bones and split digits were instead detected in *Inv1* hands. **e,** Comparison between wild-type and homozygous *Inv1* forelimbs and hindlimbs highlight the presence of underdeveloped limbs in *Inv1* as a consequence of *Fgf8* loss of expression in the AER.

Next, we went on testing whether the ectopic expression of *Btrc* and *Lbx1* in the AER of the *Dup* and *Inv1* embryos was solely due to repositioning of the *Fgf8* AER enhancers. To do so, we used CRISPR/Cas9 technology to insert four published essential *Fgf8* AER enhancers (CE58, CE59, CE61, CE66)25,26 in the *Lbx1* TAD in a wild-type background (*Fgf8 AER enh KI*, **Fig. 5a**). Given the presence of a CTCF binding site within enhancer CE58 and our aim to not create any new boundary upon knock-in we removed the CTCF recognition motif from enhancer CE58. WISH on *Fgf8 AER- enhancer KI* E11.5 mutant embryos showed misexpression of *Lbx1* and *Btrc* in the AER (**Fig. 5a**), supporting the ability of the *Fgf8* enhancers to activate other genes. Of note, we observed that *Lbx1* was ectopically expressed in the AER of all analysed embryos, while *Btrc* was present in the AER of 50% of the examined embryos and displayed a much lower signal. This result further confirmed that the observed ectopic expression of *Lbx1* and *Btrc* in the AER was due to activation by *Fgf8* enhancers.

### *Lbx1* expression in the AER induces myogenic activation

We and others have previously shown that ectopic gene expression during organ formation can lead to developmental malformation^3,4,27–29^. To investigate the molecular consequences of the high expression of *Lbx1* in the developing AER, we calculated the effect of the duplication on gene expression exclusively in the AER population, using the scRNA-seq data. We reasoned that *Lbx1* could activate its gene regulatory network in the AER and to investigate this we performed a Gene Ontology (GO) search using the 200 genes most differentially up-regulated in the AER of the *Dup* mutant compared to wild-type (**Supplementary Table 1**). Strikingly, using the MGI mammalian phenotype terms, 4 out of the 5 most enriched terms were related to myogenic phenotypes (**Fig. 5b**). Genes associated with these terms included *Lbx1*, *Myog, Gab1, Rela, Mapk14* and *Dnm2* which were all expressed in an increased number of cells in the AER of the *Dup* and at a higher level compared to wild-type (**Fig. 5c**). Of note, *Lbx1* expression was detectable in almost all AER cells while none of the wild-type AER cells expressed *Lbx1*, corroborating our previous observations. This suggested that *Lbx1* ectopic expression in the AER of the *Dup* mutant induced genes of the myogenic pathway. Additionally, expression of one of the genes, *Myog*, was also significantly increased in the bulk RNA-seq differential analysis data from limb bud in *Inv1* mutant (**Supplementary Table 2**).

### *Inv1* but not *Dup* results in a SHFM-like phenotype

Next, we analysed the mutant mice for skeletal phenotypes at developmental stage E18.5. In the *Dup* mutant we did not observe any skeletal malformations in the extremities of all the analysed transgenic embryos (n=5) nor in >25 newborns and adult mice of the established *Dup* mouse line. In contrast, 3 out of 11 E18.5 *Inv1* mutant embryos exhibited skeletal defects with a tendency to digit separation and partially missing or fused digits/bones (**Fig. 5d**), resembling the phenotype of SHFM patients13 and particularly that of the SHFM3 inversion individual (**Supplementary Fig. 2**). Indeed, clinical variability has been observed within SHFM families, between different families and also between limbs of a single individual^13,30^. Additionally, in *Inv1* we found that forelimbs and hindlimbs were underdeveloped (**Fig. 5e**), a phenotype also observed in the *Fgf8* conditional knock-out^24,31^. Loss of *Fgf8* expression is known to affect the proper development of humerus, radius, ulna, and to cause loss mainly of the first digit^24^, a phenotype different from SHFM.

To further investigate the contribution of *Lbx1* and *Btrc* to SHFM, we engineered two more CRISPR/Cas9 alleles to promote *Lbx1* or *Btrc* misexpression alone in the AER and compared it to *Dup* and *Inv1* where genes were simultaneously misexpressed. First, we generated a second, slightly larger inversion (*Inv2*) including the *Fgf8* AER enhancers and *Btrc* (**Supplementary Fig. 4b**). This mutant showed ectopic interactions between the *Fgf8* AER enhancers and *Lbx1* only, as observed in cHi-C and particularly in the subtraction cHi-C map (**Supplementary Fig. 4c and 4d**) but in general with much lower intensity compared to *Inv1* mutant. Misexpression of only *Lbx1* but not *Btrc* was indeed detected in the AER (**Supplementary Fig. 4a**). We also noticed that the embryos exhibited underdeveloped head structure most likely as a consequence of the midbrain-hindbrain boundary *Fgf8* enhancers25,26 being involved into this larger inversion (**Supplementary Fig. 4a**). To investigate the effect of an ectopic expression of *Btrc* alone, we re-targeted the *Inv1* allele and deleted *Lbx1* (*Inv1 ΔLbx1*) (**Supplementary Fig. 4b**). As expected, no misexpression for *Lbx1* was detected in the AER and in the limb in general, while *Btrc* was transcriptionally active (**Supplementary Fig. 4a**). In addition, we observed a weak expression of *Fgf8* rescued in the AER, underlining the ability of the *Fgf8* AER enhancers to re-contact their endogenous target gene in the absence of *Lbx1* and, as it happened in the *Dup* mutant, to overcome the boundary. Skeletal analysis of *Inv2* and *Inv1 ΔLbx1* mutants (**Supplementary Fig. 4f**) revealed underdeveloped limbs due to a complete (*Inv2*) or strong (*Inv1 ΔLbx1*) loss of *Fgf8* expression, but none of them (n=10 for *Inv2* and n=7 for *Inv1 ΔLbx1*) showed the level of SHFM phenotype observed in *Inv1*, supporting the idea that misexpression of both *Lbx1* and *Btrc* is required to develop a SHFM-like phenotype. Comparison between *Inv1*, *Inv2* and *Inv1 ΔLbx1* further brought up a potential effect of the partial or complete loss of *Fgf8* expression in triggering the onset of the SHFM-like phenotype in combination with *Lbx1* and *Btrc* misexpression. *Inv2* phenotype, where *Fgf8* expression was completely lost, seemed indeed more severe than *Inv1 ΔLbx1*, where *Fgf8* was only partially lost (**Fig. 5d, Supplementary Fig. 4a, 4f and Fig. 6**).

**Fig. 6.**
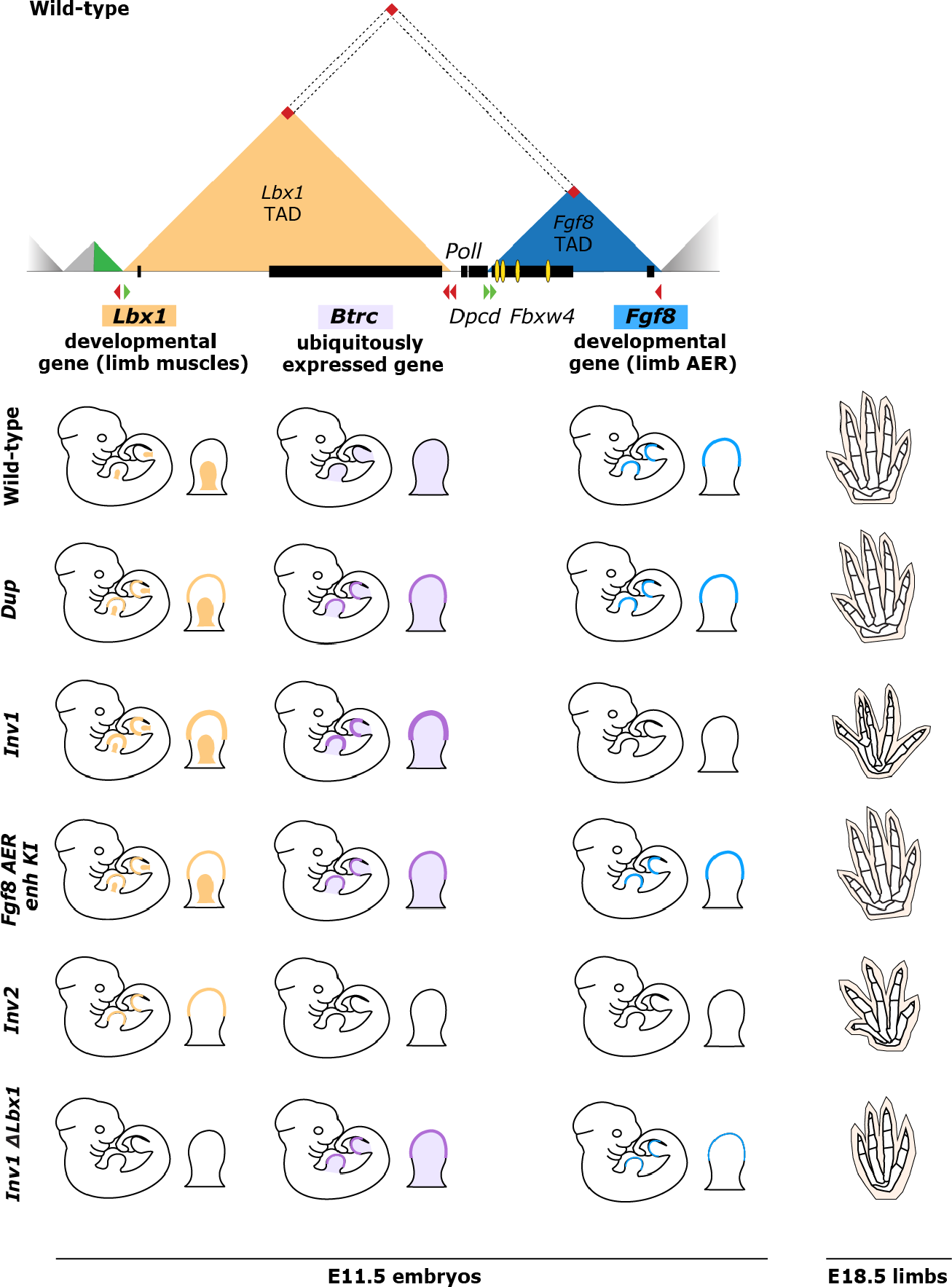
Combinatorial effects on gene expression at the *Lbx1*/*Fgf8* locus leads to SHFM3 digit fusion phenotype. Schematic overview of the *Lbx1-Fgf8* TADs (top) and *Lbx1*, *Btrc* and *Fgf8* expression in the developing limb at E11.5 and skeletal phenotype at E18.5 among the different structural rearrangements (bottom). Yellow and violet patterns represent endogenous expression in the limb bud and misexpression in the AER of *Lbx1* and *Btrc*, respectively. Turquoise pattern highlights *Fgf8* expression in the AER. Ectopic expression of both *Lbx1* and *Btrc* in the AER is observed for the *Dup*, *Inv1* and *Fgf8* AER enhancer KI (*Fgf8 AER enh KI*) mutant embryos at E11.5. However, only the *Inv1* mutants, for which the level of *Lbx1* and *Btrc* expression in the AER is higher and *Fgf8* expression is missing, show a digit fusion phenotype at E18.5, reminiscent of human SHFM. Ectopic expression of either *Lbx1* alone (*Inv2* mutant) or *Btrc* alone (*Inv1 ΔLbx1* mutant) together with complete (*Inv2*) or partial (*Inv1 ΔLbx1*) loss of *Fgf8* AER expression is not enough to lead to the digit fusion SHFM-like phenotype but only results in an *Fgf8*-KO limb phenotype.

Finally, the level of misexpression of *Lbx1* and *Btrc* played probably also an important role in developing or not the limb malformation. This could be also particularly true for the *Dup* and *Fgf8 AER-enhancer KI* mutants, where no striking phenotype was observed. Despite simultaneous misexpression of both *Lbx1* and *Btrc* genes in the AER, the level of *Btrc* misexpression was lower than in the *Inv1* mutant (**Fig. 5a and 5d**).

## Discussion

Split-hand/foot malformation (SHFM) is a congenital limb malformation affecting the central rays of the hands and/or feet (OMIM #183600, #313350, #600095, #605289, and #606708). The condition is a classic example of allelic heterogeneity, phenotypic variability, and pleiotropy^32^. cHi-C of the *Lbx1*/*Fgf8* locus, involved in SVs associated with SHFM type ^3^, showed that the region consists of two major TADs, each harbouring an important developmental gene with strikingly different expression patterns (**Fig. 6**). *Fgf8* is contained in the *Fgf8* TAD together with a number of well characterized enhancers that regulate *Fgf8* specific expression in the limb AER and other tissues^25^. *Lbx1*, a muscle-specific transcription factor expressed only in migrating muscle cells, is for its part comprised in the *Lbx1* TAD. As expected, the two regulatory TADs are separated by a boundary with divergent CTCF sites. A combination of cHi-C and 4C analysis demonstrated that the two domains are connected *via* a larger loop, established between the centromeric *Lbx1* TAD boundary and the telomeric *Fgf8* TAD boundary, within which a certain degree of contact is maintained (**Fig. 1a and Fig. 3**). This complex arrangement was disrupted by the SHFM3 duplication (*Dup*) resulting in changes in the 3D chromatin architecture. *Dup* led to ectopic interactions between the *Fgf8* AER enhancers and two other genes in the locus, *Lbx1* and *Btrc*, resulting in their misexpression in the AER in a typical *Fgf8* pattern. In contrast, *Fgf8* expression remained unchanged. This was due to the fact that *Fgf8* was not included in the duplication and a fully intact *Fgf8* regulatory landscape remained, ensuring normal expression. The duplication resulted in the formation of a neo-TAD which consisted of the truncated *Fgf8* TAD and a small region upstream of the centromeric *Lbx1* TAD boundary. As a consequence, this neo-TAD contained only *Fbwx4* with the AER enhancers and would thus not be expected to give rise to any phenotypic consequences. However, the *Fgf8* enhancers within the neo-TAD were now devoid of their target gene *Fgf8*. This was likely to result in increased interaction with other possible partners such as *Lbx1* and *Btrc*, potentiated by a pre-existing low frequency interaction between the domains, by a weaker TAD boundary upon rearrangement and also by the shorter distance particularly with *Lbx1*.

In a screen for structural variations in a cohort of patients with limb malformations, we also identified an inversion at the SHFM3 locus. Our data showed that the inversion (*Inv1*) resulted in a shift of regulatory elements from the *Fgf8* TAD into the *Lbx1* TAD thereby giving rise to the equivalent regulatory effect as the SHFM duplications. Interestingly, as for the duplication, the inversion involved the four essential *Fgf8* AER enhancers (CE58, CE59, CE61, CE66) that are located in the introns of the *Fbxw4* gene (**Fig. 1a and 1b**). Previous experiments demonstrated that a loss of *Fgf8* expression upon deletion of the four distal enhancers resulted in similar defects as conditional ablation of *Fgf8* in the limb^24,25^. In *Inv1* we also observed a loss of *Fgf8* expression and the corresponding *Fgf8* loss of function phenotype. This time, however, the enhancer region was intact but relocated from the *Fgf8* TAD into the *Lbx1* TAD. Thus, a repositioning of the boundary was sufficient to disconnect *Fgf8* from its enhancers resulting in a loss of *Fgf8* expression in the limb. The effect of *Inv1* was thus similar to the reported inversion at the *Sox9/Kcnj2* locus which resulted in a loss of *Sox9* expression due to a swap of *Sox9* regulatory elements from the *Sox9* TAD to the *Kcnj2* TAD^33^. In both SHFM-associated rearrangements, only *Lbx1* and *Btrc* were misexpressed in the AER. *Poll*, *Dpcd* and *Fbxw4* showed very low levels of expression in these cells. This indicated that these genes were not involved in SHFM3.

Our interpretation that *Lbx1* and *Btrc* were ectopically activated by *Fgf8* AER enhancers was further supported by our AER enhancers insertion experiment within the *Lbx1* TAD (*Fgf8 AER enh KI)*. As in the duplication and in the inversion, the *Fgf8* AER enhancers were able to activate both *Btrc* and *Lbx1*, with the latter being more affected. The compatibility between *Lbx1* and the *Fgf8* AER enhancers could be also explained with the fact that both *Lbx1* and *Fgf8* are developmental genes and active in the same tissue. Given a certain degree of leakiness of the *Lbx1*/*Fgf8* boundary, the contacts between the *Fgf8* AER enhancers and *Lbx1* and *Btrc* might also be expected to lead to misexpression of *Btrc* and *Lbx1* in wild-type cells. However, no cross-activation takes place. One possible explanation is that a certain threshold needs to be reached to result in gene activation and this is not achieved in the wild-type situation. In *Inv1* the level of interactions was favoured by the repositioning of the *Fgf8* AER enhancers within the *Lbx1* TAD, while in *Dup* the *Fgf8* AER enhancers in the neo-TAD lose their natural target promoter (*Fgf8*), which is not part of the neo-TAD as not included in the SV. This is likely to cause a further increase in the frequency of contacts across the leaky *Lbx1*/*Fgf8* boundary on the centromeric side as well as the newly positioned boundary telomeric of the neo-TAD, with subsequent misexpression of *Lbx1* and *Btrc*.

In *Inv1* we were able to partially recapitulate the SHFM phenotype. However, the precise patho- mechanism through which *Btrc* and *Lbx1* cause the phenotype is still unclear. The Dactylaplasia mouse mutant displays a similar phenotype to the one observed in some SHFM3 patients and the two corresponding alleles (*Dac1j* and *Dac2j*) map in the region with the duplications and inversion in SHFM334. Based on these similarities, the mutant has long been thought as a model for human SHFM3, but *Dac1j* and *Dac2j* were shown to be associated with insertions of a MusD retrotransposon upstream of the *Fbxw4* gene and within intron 5 of *Fbxw4*, respectively^34^. The absence of any structural variations in these mutants and the differences between the nature of the human and mouse genomic abnormalities argue against a common pathogenesis. However, both the mouse and the human mutations involve the regulatory elements located within and around the *Fbxw4* gene possibly providing a common pathogenetic mechanism.

Similar to the reported misexpression of *Pax3* in an *Epha4*-like pattern in the limb in brachydactyly^3^, misexpression of the muscle-specific transcription factor *Lbx1* in the AER is likely to have effects on gene expression and thus the functionality of the AER. This interpretation is supported by our finding that *Lbx1* misexpression results in the activation of muscle genes/pathways in the AER cells. At the same time, we cannot exclude a potential contribution of a loss of *Fgf8* expression to the *Inv1* mutant phenotype. The AER is the major signalling center for proximodistal growth and distal limb development. It is induced and maintained through the reciprocal interactions between the ectoderm and the underlying mesenchyme involving *Wnt-Bmp-Fgf* signaling pathways^35^. Disruption or malfunction of the AER results in diminished growth and thus hypoplasia/aplasia of digits, thus correlating with the SHFM phenotype^36^. Our results support the concept that SHFM is caused by AER defects, and in SHFM3 by gene misexpression. Mice are commonly used as a model organism to study human disease because of the similarities in genetics, physiology and organ development. However, inter-species differences exist and account for genotype-to-phenotype divergences between mouse and human^37^. When comparing CTCF binding sites in mouse and human at the *FGF8* locus, we noticed that the centromeric *Lbx1* TAD boundary is missing one binding site in reverse orientation in human in comparison to mouse (black arrow in **Fig. 3a**). This additional binding site in mouse could thus interfere with the levels of ectopic interactions between the *Fgf8* AER enhancers and *Lbx1* and *Btrc* in the *Dup* mutant, consequently reducing the level of misexpression in the AER. Indeed, gene dosage effects related to the rearrangements may explain the absence of a striking phenotype in the *Dup* in comparison to *Inv1* which is associated with highest levels of *Lbx1* and *Btrc* misexpression, causing a mouse phenotype similar to the human phenotype.

Overall, this study offers new insights into: i) the molecular phenotype of SHFM3-associated duplications and inversion, ii) gene regulation at the *Lbx1*/*Fgf8* locus in the context of 3D genome architecture and, iii) the consequences deriving from perturbations of the local chromatin structure. Notably, we provide a new and complex scenario by which SVs can cause disease. Indeed, we report in this study an example of SVs causing disease through a combined position effect mechanism resulting in ectopic gene misexpression involving multiple genes and a dose-dependent effect. To the best of our knowledge, the SVs-associated pathogenic mechanism reported in this study has not been shown previously.

In the era of whole genome sequencing (WGS) and the increased number of detected SVs, the medical interpretation of SVs and the prediction of their phenotypic consequences remain unsatisfactory. Thus, this study provides a conceptual framework when interpreting the pathogenic potential of these variant types.

## MATERIAL AND METHODS

### Mouse embryonic stem cell targeting

Culture and genome editing of mouse embryonic stem cells (mESCs) was performed as described previously^21,33^. The size and position of the human structural variations (hg19) were converted to the mouse genome (mm9) using the UCSC liftOver tool. All clones were genotyped by PCR and qPCR analyses. Positive clones were expanded, their genotyping further confirmed by PCR, qPCR and Sanger sequencing of PCR products, and used for further experiments only if the successful modification could be verified. A list of single guide RNAs (sgRNAs) used for CRISPR/Cas9 genome editing and of all the primers used for genotyping is given in **Supplementary Table 3**. For the *Fgf8-AER-enh-KI* mutant, a 6993bp DNA fragment containing the 4 *Fgf8*-AER enhancers with a deletion of the CTCF site in enhancer CE58 was synthetised and subsequently cloned into a pUC vector by *Genewiz*. Homology arms targeting the *Lbx1* locus were then cloned using Gibson assembly and the resulting vector was transfected in mESCs together with the pX459-sgRNA for CRISPR/Cas9 homology-directed-repair (HDR).

### Generation of mice

Mice were generated from genome-edited mESCs by diploid or tetraploid aggregation^38^ and genotyped by PCR and qPCR analysis. All animal procedures were conducted as approved by the local authorities (LAGeSo Berlin) under license numbers G0243/18 and G0176/19.

### Whole mount *in situ* hybridization (WISH)

RNA expression in E11.5 mouse embryos from wild-type and mutants was assessed by WISH using digoxigenin (DIG)-labelled antisense riboprobes for *Lbx1*, *Btrc*, *Poll*, *Dpcd*, *Fbxw4* and *Fgf8* transcribed from linearized gene-specific probes (PCR DIG Probe Synthesis Kit; Roche). Primers for probe generation are listed in **Supplementary Table 3**. Embryos were collected and fixed overnight in 4% PFA in PBS, then washed twice for 30 min in PBS with 0.1% Tween (PBST), dehydrated for 30 min each in 25%, 50% and 75% methanol in PBST and stored at −20 °C in 100% methanol. For WISH, genotyped embryos were rehydrated on ice in reverse methanol/PBST steps, washed twice in PBST, bleached in 6% H2O2 in PBST for 1 h and washed in PBST. Embryos were washed in PBST and refixed for 20 min with 4% PFA in PBS, 0.2% glutaraldehyde and 0.1% Tween. After further washing steps with PBST, embryos were incubated at 68 °C in L1 buffer (50% deionized formamide, 5× SSC, 1% SDS, 0.1% Tween 20 in diethyl pyrocarbonate (DEPC), pH 4.5) for 10 min. For prehybridization, embryos were incubated for 2 h at 68 °C in hybridization buffer 1 (L1 with 0.1% tRNA and 0.05% heparin), while for subsequent probe hybridization, embryos were incubated overnight at 68 °C in hybridization buffer 2 (hybridization buffer 1 with 0.1% tRNA, 0.05% heparin and 1:500 DIG probe). The next day, unbound probes were washed away through repeated washing steps: 3 × 30 min at 68 °C with L1, L2 (50% deionized formamide, 2× SSC, pH 4.5, 0.1% Tween 20 in DEPC, pH 4.5) and L3 (2× SSC, pH 4.5, 0.1% Tween 20 in DEPC, pH 4.5). For signal detection, embryos were treated for 1 h with RNase solution (0.1 M NaCl, 0.01 M Tris, pH 7.5, 0.2% Tween 20, 100 μg ml−1 RNase A in H2O), followed by washing in TBST 1 (140 mM NaCl, 2.7 mM KCl, 25 mM Tris–HCl, 1% Tween 20, pH 7.5) and blocking for 2 h at room temperature in blocking solution (TBST 1 with 2% calf serum and 0.2% bovine serum albumin). Overnight incubation with Anti-Dig antibody conjugated to alkaline phosphatase (1:5,000) at 4 °C (no. 11093274910; Roche) was followed by 8 × 30 min washing steps at room temperature with TBST 2 (TBST with 0.1% Tween 20 and 0.05% levamisole–tetramisole) and left overnight at 4 °C. Embryos were finally stained after equilibration in AP buffer (0.02 M NaCl, 0.05 M MgCl2, 0.1% Tween 20, 0.1 M Tris–HCl and 0.05% levamisole–tetramisole in H2O) 3 × 20 min, followed by staining with BM Purple AP Substrate (Roche). The stained embryos were imaged using a Zeiss SteREO Discovery V12 microscope and Leica DFC420 digital camera.

### Skeletal preparation

E18.5 fetuses were kept in H2O for 1–2 h at room temperature and heat shocked at 65 °C for 1 min. The skin was gently removed, together with the abdominal and thoracic viscera using forceps. The fetuses were then fixed in 100% ethanol overnight. Afterwards, alcian blue staining solution (150 mg l−1 alcian blue 8GX in 80% ethanol and 20% acetic acid) was used to stain the cartilage overnight. After 24 h fetuses were rinsed and postfixed in 100% ethanol overnight, followed by 24 h incubation in 0.2% KOH in H2O for initial clearing. The next day fetuses were incubated in alizarin red (50 mg l−1 alizarin red S in 0.2% potassium hydroxide) to stain the bones overnight. Following this, rinsing and clearing was done for several days using 0.2% KOH. The stained embryos were dissected in 25% glycerol, imaged using a Zeiss SteREO Discovery V12 microscope and Leica DFC420 digital camera, and subsequently stored in 80% glycerol.

### Capture Hi-C

E11.5 mouse limb buds were prepared in 1x PBS and dissociated by trypsin treatment for 10 min at 37°C, pipetting every 2 minutes to obtain a single-cell suspension. Trypsin was stopped by adding 5x volume of 10% FCS/PBS. The solution was then filtered through 40 μm cell strainer (Falcon) to remove cell debris, cells were centrifuged at 1100 rpm for 5 min and the pellet was then resuspended in 5 ml 10% FCS/ PBS. 5 ml of freshly prepared 4% formaldehyde in 10% FCS/PBS (final concentration 2%) were added to perform crosslinking and samples were incubated in rotation for 10 min at room temperature. The crosslinking reaction was stopped by adding 1 ml 1.425M glycine on ice, followed by centrifugation at 1500 rpm and 4°C for 8 min, pellet resuspension in 5 ml freshly prepared, cold lysis buffer (final concentration of 10 mM Tris pH 7.5, 10 mM NaCl, 5 mM MgCl2, 0.1 M EDTA, and 1 × cOmpleteTM protease inhibitors (Sigma-Aldrich) and incubation for at least 10 min on ice. Nuclei were pelleted by centrifugation at 2000 rpm at 4°C for 5min, washed with 1x PBS, aliquoted in 1.5ml tubes with 2.5-5 x 106 nuclei each, followed by supernatant removal, snap-freezing and storage at -80°C.

Snap-frozen pellet was resuspended in 360 μl of water, mixed with 60 μl of 10x DpnII buffer (Thermo Fisher Scientific) and incubated for 5 min at 37°C in a thermomixer at 900 rpm. 15 μl of 10% SDS was then added and the samples were incubated for 1 h at 37°C and 900rpm, with occasional pipetting to help dissolving the nuclei aggregates. Next, 150μl of 10% Triton X-100 were added, followed by 1 h of incubation at 37°C and 900rpm. After adding 600 μl of 1x DpnII buffer, a 10 μl aliquot was taken as undigested control and stored at 4°C and 400 units of restriction enzyme DpnII were added to the samples. After 4 h of digestion additional 200 units of DpnII enzyme were added and the samples were incubated overnight at 37°C with shaking at 900 rpm. After overnight incubation, a 10 μl aliquot was taken as digested control and stored at 4°C, while the samples were supplemented with further 200 units of DpnII enzyme for additional four hours. Meanwhile, the undigested and digested controls were tested. For this, each 10 μl aliquot was mixed with 85 μl 10mM Tris pH 7.5 and 2 μl RNase A (10 mg/ml) and incubated for 1 h at 37°C. Chromatin was decrosslinked by adding 5 μl proteinase K (10 mg/ ml) and incubation at 65°C for 4 h. The DNA was then extracted by adding 100 μl phenol-chloroform. Samples were mixed by inverting the tubes and centrifuged for 10 min at 13200 rpm at room temperature. The upper water phase was transferred into new 1.5ml tubes and analysed on a 1% agarose gel. After correct validation, restriction enzyme in the original samples still under digestion was heat-inactivated at 65°C for 20 min. The samples were transferred to 50 ml Falcon tubes and 700 μl 10x ligation buffer (Thermo Fisher Scientific) were added. The volume was filled up to 7 ml with water and 50 units of T4 DNA ligase (Thermo Fisher Scientific) were added. The ligation mix was incubated overnight at 4°C in rotation. The next day, a 100 μl aliquot of religated DNA was collected and analysed on an agarose gel as described above. Upon successful ligation, samples were de-crosslinked by adding 30 μl of proteinase K (10 mg/ml) and incubating overnight at 65°C. Then, 30 μl RNase A (10 mg/ml) were added and the samples were incubated for 45 min at 37°C. The DNA was then extracted by adding 7 ml phenol-chloroform. The solution was mixed by inverting the tube and the water phase was separated by centrifugation at 3750 rpm for 15 min at room temperature. DNA was precipitated by adding the following reagents to the water phase: 7 ml water, 1.5 ml 2M NaAc pH 5.^6^, 140 μg glycogen, 35 ml 100 % ethanol. The solution was mixed and placed at -80°C overnight or until the samples were completely frozen. The sample was then thawed and centrifuged for20 min at 8350 g and 4°C. DNA pellet was washed with 30 ml cold 70 % ethanol and centrifuged 15 min at 3300 g at 4°C. Dried pellet was dissolved in 150 μl 10mM Tris pH 7.5 at 37°C. The final 3C library was checked on 1% agarose gel and subsequently used for capture Hi-C preparation. 3C libraries were sheared using a Covaris sonicator (duty cycle: 10%; intensity: 5; cycles per burst: 200; time: 6 cycles of 60 s each; set mode: frequency sweeping; temperature: 4–7 °C). Adaptors were added to the sheared DNA and amplified according to Agilent instructions for Illumina sequencing. The library was hybridized to the custom-designed SureSelect beads and indexed for sequencing following Agilent instructions. SureSelect enrichment probes were designed over the genomic interval chr19:44,440,000-46,400,000 (mm9) using the online tool of Agilent: SureDesign. Probes were covering the entire genomic region and were not designed specifically in proximity of DpnII sites. All samples were then sequenced with the Hiseq4000 or Novaseq6000 Illumina technology according to the standard protocols and with around 400 million 50bp, 75bp (Hiseq4000) or 100bp (Novaseq6000) paired-end reads per sample.

### Capture Hi-C data processing

In a pre-processing step, the reads in fastq files were trimmed to 50 bp, if necessary, to obtain the same initial read length for all samples. Afterwards, mapping, filtering and deduplication of paired-end sequencing data was performed using the HiCUP pipeline v0.8.1^39^ (no size selection, Nofill: 1, Format: Sanger). The pipeline was set up with Bowtie2 v2.4.2^40^ for mapping short reads to reference genome mm^9^. For merging biological replicates, the final bam files produced by the HiCUP pipeline were joined. Juicer tools v1.19.02^41^ was used to generate binned and KR normalized^42,43^ contact maps from valid and deduplicated read pairs. For the generation of cHi-C maps, only read-pairs referring to the region of interest (chr19:44,440,001-46,400,000) and with MAPQ≥30 were considered. We used cHi-C maps with 5kb bin size. For the sample with a duplication it is noted, that the matrix balancing of the KR normalization affects the signal in the duplicated region, because it scales down the signal intensities in this region to fit to the other regions of the cHi-C map. In order to consider the duplicated region explicitly, we created for the *Dup* mutant sample an additional version of the cHi-C map by applying LOIC normalization^44^ (python package iced v.0.5.1045) to raw count map, which has the aim to retain the effects of the increased copy number. For the LOIC normalization the copy number was set to 4 for all bins overlapping the duplication, and to 2 for the other bins. For five matrix rows/columns with low coverage, the count values were removed prior to LOIC-normalization.

Subtraction maps were computed from KR-normalized cHi-C maps, which were normalized in a pairwise manner before subtraction as follows. To account for differences between two maps in their distance-dependent signal decay, the maps were scaled jointly across their sub-diagonals. Therefore, the values of each sub-diagonal of one map were divided by the sum of this sub-diagonal and multiplied by the average of these sums from both maps. Afterwards, the maps were scaled by 106/total sum. For the computation of scaling factors, the duplicated as well as the inverted regions were excluded in both maps. cHi-C maps were visualized as heatmaps with linear scale, with very high values being truncated to improve visualization.

### Virtual 4C

Virtual 4C-like interaction profiles were generated for individual viewpoints from the same bam files also used for the cHi-C maps. Paired-end reads with MAPQ≥30 were considered in a profile, when one read mapped to the defined viewpoint region and the other one outside of it. Contacts of a viewpoint region were counted per restriction fragment. The count profile was binned to a 1 kb grid. In case a fragment overlapped more than one bin, the counts were distributed proportionally. Afterwards, the binned profile was smoothed by averaging within a sliding window of 5kb size and scaled by 103/sum of counts within the enriched region. The viewpoint region itself and a margin ±5kb around it were excluded from the computation of the scaling factor. Interaction profiles were generated with custom Java code using v2.12.0 (https://samtools.github.io/htsjdk/).

### RNA-seq

E11.5 limb buds were microdissected from wild-type and mutant embryos (*n* = 3) and immediately frozen in liquid nitrogen. Total RNA was isolated using the RNeasy Mini Kit (QIAGEN). Samples were poly-A enriched and sequenced with the Novaseq6000 Illumina technology according to the standard protocols and with around 100 million 100bp paired-end reads per sample.

### RNA-seq data processing

Reads were mapped to the mouse reference genome (mm9) using the STAR mapper^46^ (splice junctions based on RefSeq; options: –alignIntronMin 20–alignIntronMax 500000– outFilterMultimapNmax 5–outFilterMismatchNmax 10–outFilterMismatchNoverLmax 0.1). Reads per gene were counted as described previously^3^, and used for differential expression analysis with the DEseq2 package^47^.

### scRNA-seq

scRNA-seq experiments were performed in single replicates that were jointly processed to avoid batch effects. E11.5 hindlimb buds of wild-type and Dup embryos were microdissected in 1xPBS at room temperature. A single-cell suspension was obtained by incubating the tissue for 10 min at 37 °C in 200 μl Gibco trypsin-EDTA 0.05% (Thermo Fisher Scientific) supplemented with 20 μl 5% BSA (Sigma-Aldrich). Trypsinization was then stopped by adding 400 μl of 5% BSA. Cells were then resuspended by pipetting, filtered using a 0.40 μm cell strainer (Falcon), washed once with 0.04% BSA, centrifuged for 5 min at 1200 rpm, then resuspended in 0.04% BSA. The cell concentration was determined using an automated cell counter (Bio-Rad) and cells were subjected to scRNA-seq (10x Genomics, Chromium Single Cell 3ʹ v2; one reaction per time point and per strain has been used) aiming for a target cell recovery of up to 10,000 sequenced cells per sequencing library (time point and strain). Single-cell libraries were generated according to the 10x Genomics instructions with the following conditions: 8 PCR cycles were run during cDNA amplification and 12 PCR cycles were run during library generation and sample indexing to increase library complexity. Libraries were sequenced with a minimum of 230 million 75bp paired-end reads according to standard protocols.

### scRNA-seq data processing

The Cell Ranger pipeline v.3 (10x Genomics lnc.) was used for each scRNA-seq sample in order to de-multiplex the raw base call files, to generate the fastq files, and to perform the alignment against a custom mouse reference genome mm9 to create the unique molecular identifier (UMI) count matrix. Only cells with more than 2000 detected genes and less than 10% of mitochondrial UMI counts were considered for further analysis. In addition, Scrublet^48^ was used to identify potential doublets in our dataset. Cells with a Scrublet score higher than 0.2 were filtered out. Each sample was normalized independently using the SCT method^49^ implemented in Seurat3^50^ R package and then, these were integrated using the Seurat3 IntegrateData CCA-based (canonical correlation analysis) approach considering the top 2,000 most variable genes. The first 20 principal components of this joint dataset were calculated and used for UMAP projection and reconstruction of the cell-cell similarity graph. To delimitate the major hindlimb bud cell types, we used the Louvain algorithm implemented in the Seurat3 function FindClusters and the expression of well-known marker genes for the limb-comprising cell types. For the AER cluster, we used the marker gene *Fgf8*. Differential gene expression between the WT and the Dup cells was estimated by modelling the gene expression as a function of the genotype across cells. We fitted a quasi-Poisson distribution to calculate the effect of the *Dup* genotype on the gene expression distribution of each gene using the monocle3 strategy^51^. We tested that such effect was not equal to 0. We use this condition association effect to rank the genes and identified the pathways associated with the top 200 genes using the Enrichr tool from the Maayan lab^52^. Finally, we confirmed that the top perturbed genes associated to the muscle pathways are more frequently expressed in the *Dup* AER cells.

## 4C-seq

For 4C-seq libraries, 5×106 – 107 cells were used. The fixation and lysis were performed as described in the Capture Hi-C section. After the first digestion with DpnII (NEB), sticky ends were religated in a 50 ml falcon tube (700 μl 10 ligation buffer (Fermentas), 7 ml H2O, 50 U T4 DNA ligase (Thermo); overnight at 16 °C) and DNA de-cross linked and cleaned as described in the Capture Hi-C section. Next, a second digestion (150 μl sample, 50 μl 10× Csp6I buffer (Thermo), 60 U Csp6I (Thermo) 295 μl H2O; overnight at 37 °C) and another re-ligation were performed. For all viewpoints, DNA was purified using a PCR clean up Kit (Qiagen) and 1.6 μg DNA was amplified by PCR (*LBX1* Viewpoint: read-primer 5ʹ-TCTATATGCTACCATGATC-3ʹ, secondary-primer 5ʹ- GATGAACTGGAATACCCA-3ʹ; *FGF8* Viewpoint: read-primer 5ʹ-AGGGTGCGTTCCAAGATC-3ʹ, secondary-primer 5ʹ-GGTGGCCTGGATGGAAGT-3ʹ; *BTRC* Viewpoint: read-primer 5ʹ- CAACGCAGCGCCCGGATC-3ʹ, secondary-primer 5ʹ-CTGGGAATGAGGACCTAGGGC-3ʹ). For the library reaction, primers were modified with TruSeq adapters (Illumina): Adapter1 5ʹ- CTACACGACGCTCTTCCGATCT-3ʹ and Adapter2 5ʹ-CAGAC GTGTGCTCTTCCGATCT-3ʹ. Between 50 and 200 ng were used as input of a single 4C PCR reaction depending on the complexity. The reaction was performed in a 50 μl volume using the Expand Long Template System (Roche) and 29 reaction cycles. After the PCR all reactions were combined and the DNA purified with a PCR clean up Kit (Qiagen). All samples were then sequenced with the Novaseq6000 Illumina technology according to the standard protocols and with around 5 million 100bp single-end reads per sample.

### 4C-seq data processing

Sequencing reads were mapped, normalized and smoothed with pipe4C^53^ using the reference genome GRCh37 and default settings. All viewpoints were performed in replicates and as quality measure >70% of reads were mapped within a size range of 1Mb and >80% within 100kb around the viewpoint.

### Human material

Skin biopsies were collected from SHFM3 patients and controls by standard procedures. Fibroblasts were cultured in DMEM (Lonza) supplemented with 10% fetal calf serum (Gibco), 1% L-glutamine (Lonza) and 1% penicillin/streptomycin (Lonza). Informed consent was obtained from all individuals that participated in this study. This study was approved by the Charité Universitätsmedizin Berlin ethics committee.

## ACKNOWLEDGEMENTS

This study was supported by a grant from the Deutsche Forschungsgemeinschaft (MU 880/22-1) to S.M. We thank the transgenic unit, sequencing core and animal facility of Max Planck Institute for Molecular Genetics for technical assistance, Ute Fischer for technical support and Norbert Brieske for help with whole mount *in situ* hybridizations and image processing. We would like to thank all members of the Mundlos laboratory and Tugce Aktas for continuous support and discussion. We would like to particularly thank Dario Lupiáñez, Chiara Anania, Lila Allou and Jane Skok for critical reading of the manuscript.

## AUTHOR CONTRIBUTIONS

G.C., S.M. and M.S. conceived the project. G.C. performed most experiments with major support from J.G and further technical support of M.F., R.F. and M.F. L.W. performed morula aggregation.

R.S. performed cHi-C computational analysis and generated virtual 4C maps. S.A. performed and analysed 4C-seq. C.A.P.-M. performed Single Cell RNA-seq computational analysis. B.T. oversaw high-throughput sequencing. M.S., E.P., R.F., F.N., M.G., and O.Z. examined the patients, interpreted the clinical data and provided the clinical information together with biological samples. G.C. and S.M. wrote the manuscript with input from all the authors.

## SUPPLEMENTARY FIGURES

**Supplementary Fig. 1.**
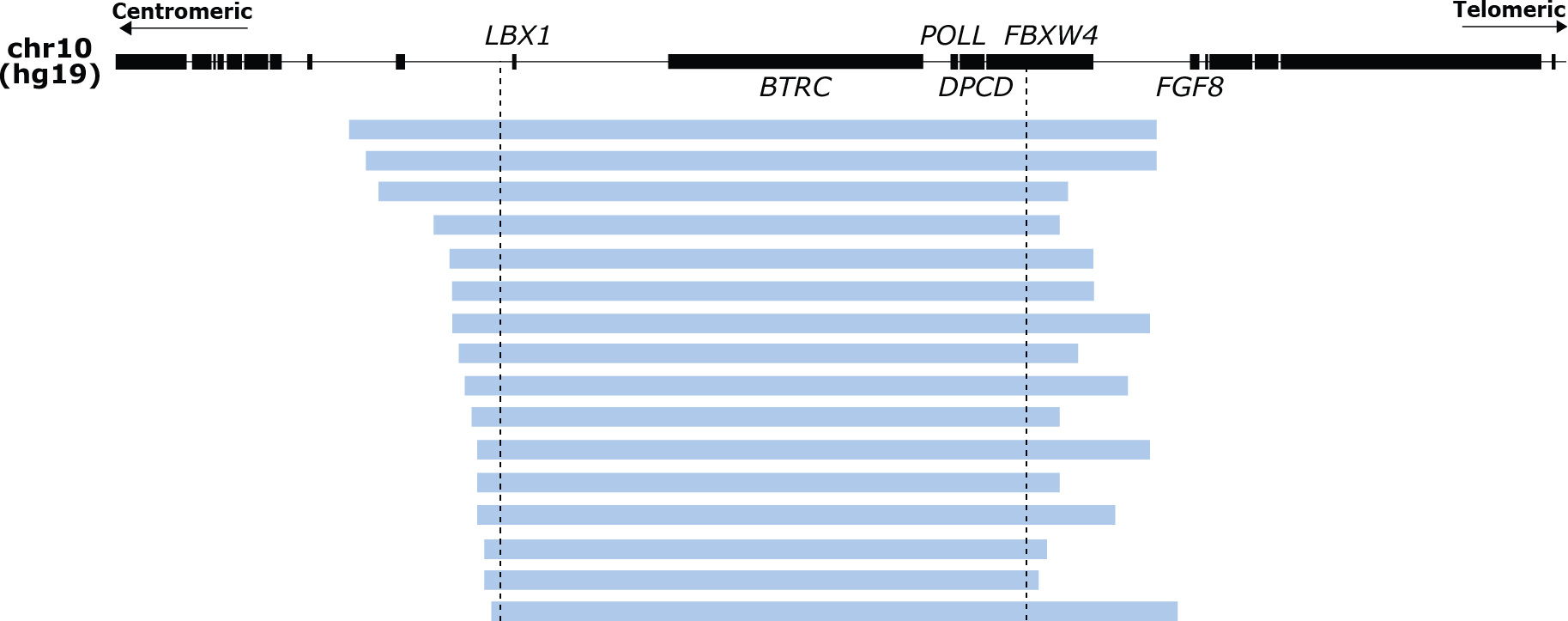
Overview of in house SHFM3 duplications. All the identified duplications included *LBX1* at the centromeric side and excluded *FGF8* at the telomeric side. Dashed black lines indicate the minimal critical region of the duplications.

**Supplementary Fig. 2.**
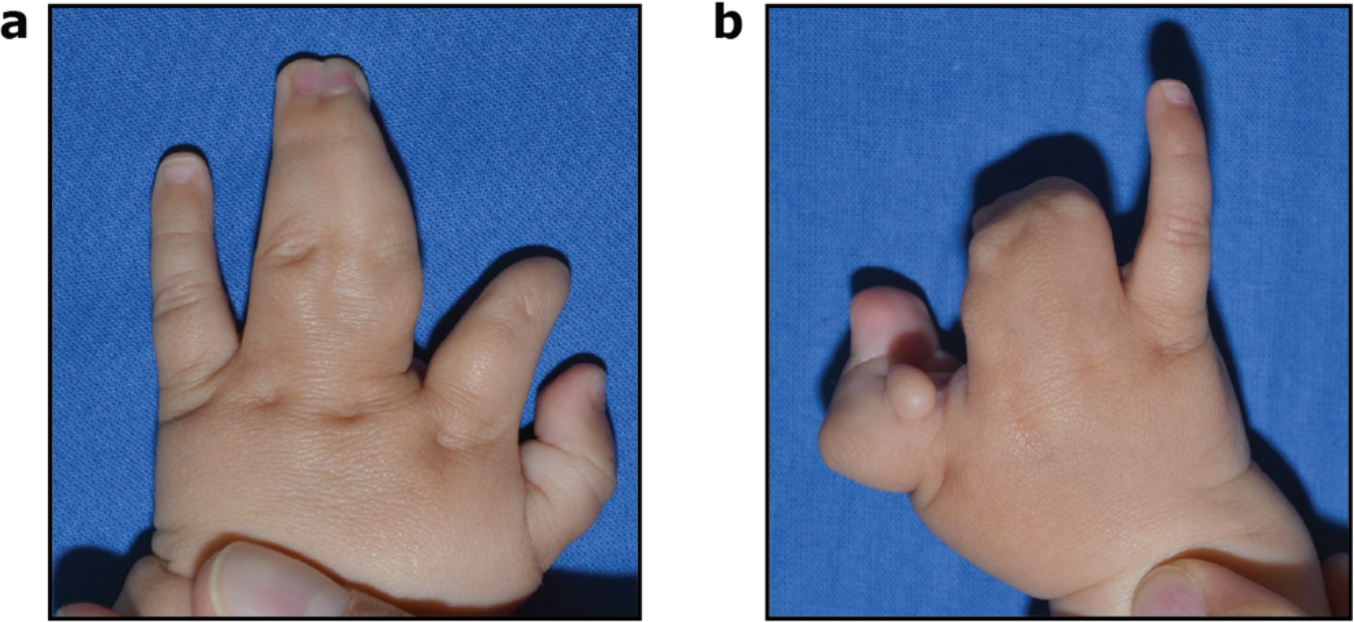
Patient carrying an inversion at the *LBX1*/*FGF8* locus exhibited two phenotypes of the SHFM spectrum. **a**, Left hand showing fusion of digit III and IV. **b,** Right hand showing absence of central distal digits.

**Supplementary Fig. 3.**
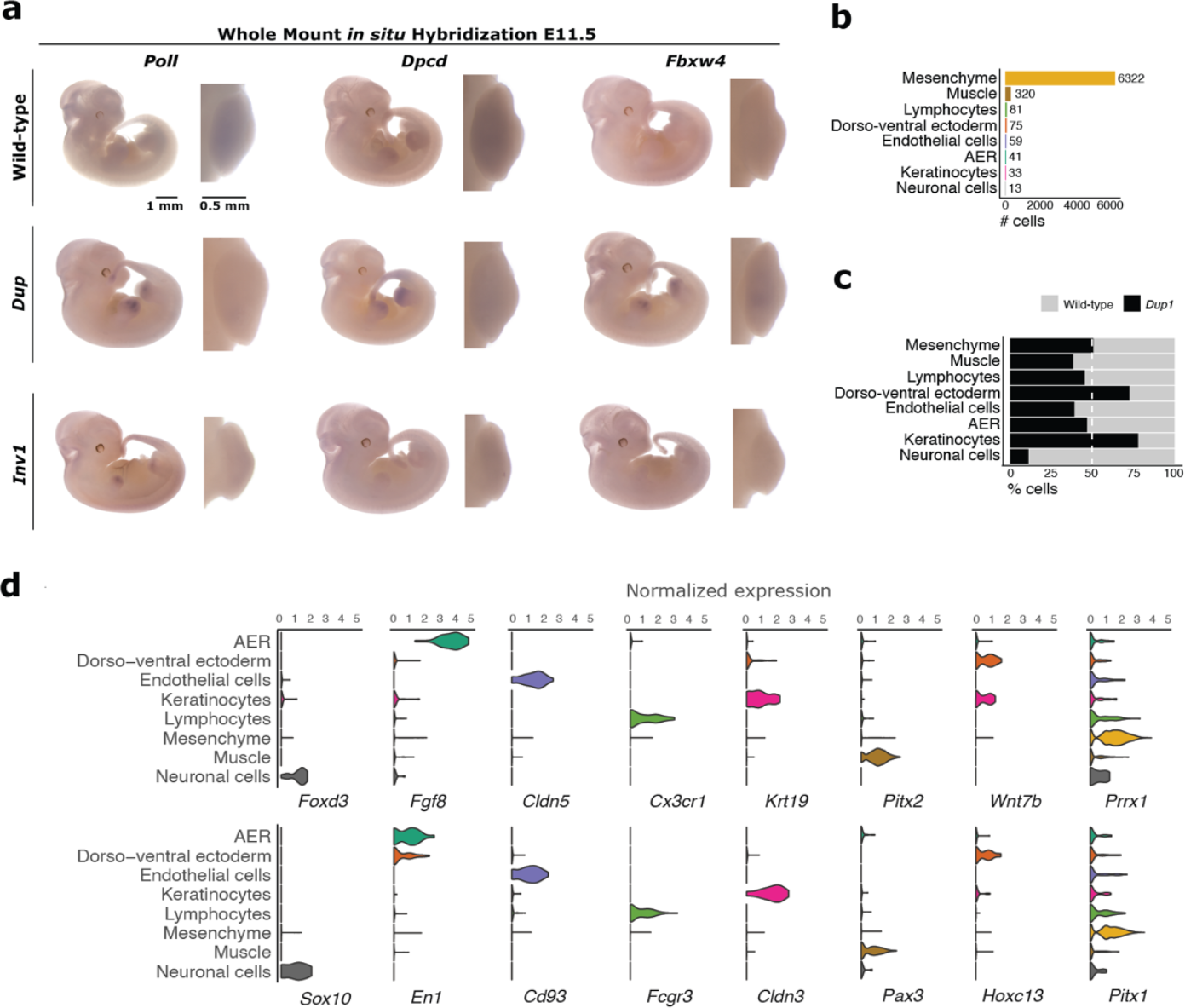
No misexpression in the AER was detected for other genes at the *Lbx1-Fgf8* locus. **a**, WISH of the other genes at the *Lbx1-Fgf8* locus for *Dup* and *Inv1*. Expression was checked and confirmed in at least 3 or more homozygous embryos. **b,** Bar plot representing the number of cells per cluster as defined in Figure 4b. These numbers correspond to the merged data of both wild-type and *Dup* samples, from the scRNA-seq of E11.5 hindlimbs. **c,** Bar plot representing the % of cells per cluster as defined in Figure 4b for each genotype separately. Some biases are observed, particularly for small cell population, as expected from scRNA-seq data. **d,** Violin plots representing the expression of two marker genes per cluster that were used to confirm the clustering analysis shown in Figure 4b. By example, AER cells are defined by high expression of *Fgf8* and *En1*.

**Supplementary Fig. 4.**
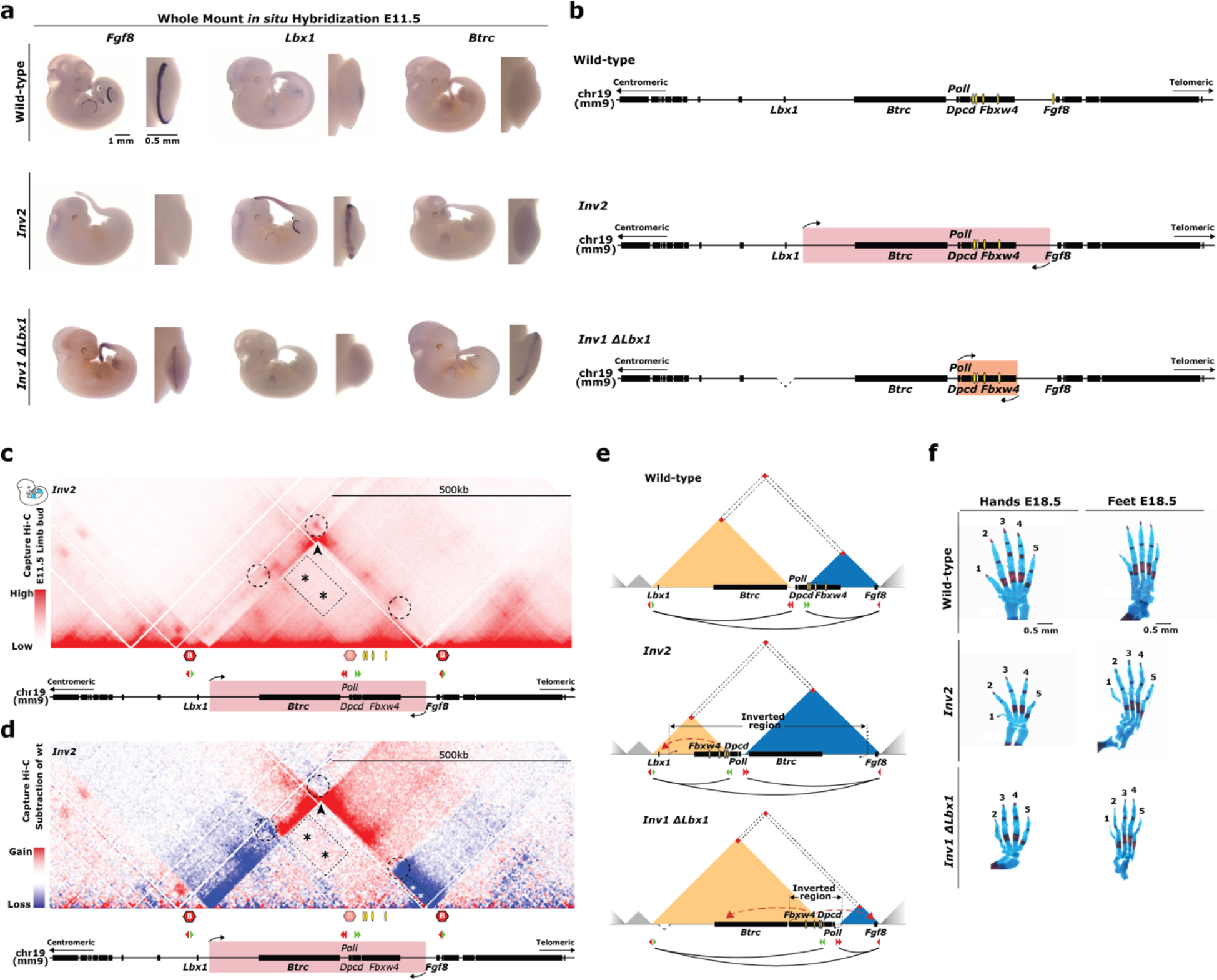
Misexpression of both *Lbx1* and *Btrc* is required to develop a SHFM phenotype. **a**, Whole-mount *in situ* hybridization for *Fgf8*, *Lbx1* and *Btrc* at E11.5. Expression was checked and confirmed in at least 3 or more homozygous embryos. **b,** Schematics of the locus for wild-type and CRISPR/Cas9 alleles. Yellow ovals highlight *Fgf8* AER enhancers. **c,** cHi-C of homozygous *Inv2*. Dashed circles indicate the position of the original wild-type interactions between boundaries, two of them lost upon reshuffling of the boundary between *Lbx1* and *Fgf8* TADs as a consequence of this inversion. The boundary involved in the inversion is shown as blurry. Black arrowhead highlights the bow tie configuration representative of the inverted regions. The rectangular dashed area highlighted by asterisks show ectopic interactions compared to wild-type between the *Fgf8* AER enhancers region and the one containing now only *Lbx1*. **d,** Subtraction of wild-type cHi-C interactions from *Inv2* mutants. Dashed circles highlight the loss (blue) of the original interactions between boundaries as a consequence of TAD- reshuffling. Black arrowhead highlights the classic bow tie configuration representative of the inverted regions. The rectangular dashed area highlighted by black asterisks show ectopic interactions compared to wild-type between the *Fgf8* AER enhancers region and the one containing *Lbx1* only, while the rectangular dashed area highlighted by white asterisks points out the loss of interactions between *Fgf8* and its AER enhancers. **e,** (Top) Wild-type schematic. (Middle) *Inv2*. (Bottom) *Inv1 ΔLbx1*. Upon *Inv2* the *Fgf8* regulatory elements are repositioned within a smaller *Lbx1* TAD without *Btrc*, so that the ectopic interactions are established only with *Lbx1* (red dashed arrow), whereas the *Fgf8* TAD is increased in size and comprises not only *Fgf8*, but also *Btrc*, both isolated from *Fgf8* enhancers. **f,** Skeletal analysis of E18.5 hands and feet stained with alcian blue (cartilage) and alizarin red (bone) from wild-type, *Inv2* and *Inv1 ΔLbx1*.

